# Detecting Inversions with PCA in the Presence of Population Structure

**DOI:** 10.1101/736900

**Authors:** Ronald J. Nowling, Krystal R. Manke, Scott J. Emrich

## Abstract

Chromosomal inversions are associated with reproductive isolation and adaptation in insects such as *Drosophila melanogaster* and the malaria vectors *Anopheles gambiae* and *Anopheles coluzzii*. While methods based on read alignment have been useful in humans for detecting inversions, these methods are less successful in insects due to long repeated sequences at the breakpoints. Alternatively, inversions can be detected using principal component analysis (PCA) of single nucleotide polymorphisms (SNPs). We apply PCA-based inversion detection to a simulated data set and real data from multiple insect species, which vary in complexity from a single inversion in samples drawn from a single population to analyzing multiple overlapping inversions occurring in closely-related species, samples of which that were generated from multiple geographic locations. We show empirically that proper analysis of these data can be challenging when multiple inversions or populations are present, and that our alternative framework is more robust in these more difficult scenarios.

## INTRODUCTION

Chromosomal inversions play an important role in ecological adaptation by enabling the accumulation of beneficial alleles (Love et al. (2016); Fuller et al. (2018); Prevosti et al. (1988)) and reproductive isolation (Noor et al. (2001)). For example, the 2La inversion in the *Anopheles gambiae* complex has been associated with thermal tolerance of larvae (Rocca et al. (2009)), enhanced desiccation resistance in adult mosquitoes (Gray et al. (2009)), and susceptibility to at least one species (*Plasmodium falciparum*) of malaria (Riehle et al. (2017)).

Inversion analysis contains three sub-problems: detection (is an inversion present?), localization of an inversion along a chromosome arm, and determining the orientations of inversions present in each sample (karyotyping). Most techniques can perform a subset of these tasks, but not all of them. For example, some insects such as *Drosophila melanogaster* and the mosquito *Anopheles gambiae* have large polytene chromosomes, which can be seen directly under a microscope. This enables detection and karyotyping of previously characterized inversions (Lobo et al. (2010); Sharakhov et al. (2006); George et al. (2010)).

Computational approaches developed for model organisms such as human – or species without visible chromosomes including many other insects – are generally based on sequencing large DNA fragments from alternative karyotypes. Specifically, inversion breakpoints relative to a known reference genome can discovered by checking for cases where either mate-pair or long-read sequence data align unexpectedly (e.g., Zhu et al. (2017); Corbett-Detig et al. (2012); Hormozdiari et al. (2009); Chen et al. (2009); Suzuki et al. (2014); Zhu et al. (2018)). Breakpoints in *Anopheles* mosquitoes are characterized by long, repeated sequences (Sharakhov et al. (2006); Lobo et al. (2010)), however, which has prohibited break point detection using these existing sequence alignment-based methods (Zhu et al. (2017, 2018)).

An alternative approach that can use single-nucleotide polymorphism (SNP) data would be even more attractive because it would not require specialized sequencing (e.g., long reads generated from high molecular weight DNA). SNP data are used for a wide range of analyses and are inexpensive to generate using commonly-available next-generation sequencing (NGS) techniques. Prior work has used Principal Component Analysis (PCA). For example, PCA of SNP data is widely used in population genetics to visualize the relationships between samples (Neafsey et al. (2010)), correcting for stratification in genome-wide association studies (Price et al. (2006)), and with clustering to determine population structure (Lee et al. (2009); Patterson et al. (2006)).

Inversion differences within a population can also appear as clusters in PCA projections (Ma and Amos (2012); Ma et al. (2014)), which has motivated computational detection based on characterizing this observed cluster structure (Cáceres and González (2015)). Because not all data induce a clear pattern in PCA projection plots, we were motivated to develop an alternative method based on single-SNP association tests (see Nowling and Emrich (2018c)). PCA is first performed on the entire set of SNPs from a single chromosome. For each PC, single-SNP association tests are performed against the samples’ projected PC coordinates. The spatial relationships of the associations are then visualized with Manhattan plots to reveal inversions. We applied this method to 34 *An. gambiae* and *An. coluzzii* samples (from Fontaine et al. (2015)) from four geographic locations. No clear cluster structure was distinguishable due to small sample sizes and confounding factors, but our method still was able to successfully detect and localize a major inversion (2La, confirmed against experimental karyotyping labels) and multiple inversions on 2R.

Here, we focus on factors that we found confound PCA-based cluster analysis. We note that prior work (see Ma and Amos (2012); Cáceres and González (2015)) focus on human genomes, which tend to be easier for making inferences. In support of this, we use invertFREGENE to simulate and evaluate an ideal situation with a single population and a single inversion. Using *Drosophila* and *Anopheles* data, however, provides test cases for evaluating large inversion detection when the biology is not as clear. For example, the 198 *Drosophila melanogaster* fly samples from the *Drosophila* Genetics Reference Panel 2 (DGRP2) (Mackay et al. (2012); Huang et al. (2014)) include multiple, overlapping inversions on the 3R chromosome arm. *Anopheles* data have been previously analyzed with PCA and found to cluster based on combinations of inversion karyotype, species, and geography (Fontaine et al. (2015); Neafsey et al. (2010); Miles et al. (2016); Nowling and Emrich (2018c)). This allows using 150 Burkina Faso *An. gambiae* and *An. coluzzii* mosquito samples to look at the effect of species–inversion interactions (Miles et al. (2016)), and the re-analysis of the 34 *An. gambiae* and *An. coluzzii* samples (from Fontaine et al. (2015)) to look more deeply at species–population–inversion interactions.

We confirm that identification and localization of inversions using PCA can be an easier task because the clustering required for karyotyping is not always clear. For example, Cáceres and González used Gaussian mixture models to cluster samples from PCA projections and then performed a likelihood-ratio test based on the presence of three clusters corresponding to the three expected inversion orientations (Cáceres and González (2015)). The clusters obtained from these well-characterized insect data with experimentally determined karyotypes, however, are not always the three expected inversion orientations. Using our framework, we then tried performing single-SNP association tests against the cluster labels (instead of against the projected PC coordinates) to determine if they are more robust. Although we could accurately infer karyotypes, we also remain susceptible to data with either multiple inversions or from closely related species. This is in some sense expected given the role of PCA in population inference and other more traditional population genetics analysis (Lee et al. (2009); Patterson et al. (2006); Price et al. (2006); Neafsey et al. (2010)). For these more complex cases, we show that populations need to be analyzed individually and care must be taken when choosing which PCs and cluster number to use. We show that our PC-SNP association tests are easier to use and more robust in large part since they do not depend on accurately clustering samples to detect inversions like other PCA-based approaches.

## METHODS

### Data Sets

We use invertFREGENE for the simulated data set (O’Reilly et al. (2010)). We use default parameters for the mutation rate (2.3×10^−7^), recombination rate (1.25×10^−7^), proportion of crossovers in recombination hot spots (0.88), length of crossover hot spots (2000), per-base gene-conversation rate (4.5 10^−8^), and gene-conversation length (500). We simulate 1000 2Mb haploid chromosomes (created from a single founder) in one population and no inversions for 10,000 generations to equilibrate. We introduce an inversion from 0.75 Mb to 1.25 Mb and continued the simulation for another 10,000 generations (or until the inversion frequency reached 50%). We set the MaxFreqOfLostInv parameter to 10% and set the output mode to “sequence” mode. We modify invertFREGENE to output inversion orientations of the haploids. We wrote a custom script in Python to randomly sample haploids without replacement to produce diploid individuals and write a VCF.

We also use three real and publicly-available data sets. For the samples from Fontaine et al. (2015), we retrieve the VCF files from the Dryad Digital Repository (Fontaine et al. (2014)), sample IDs from the supplemental materials of the paper, and use VCFtools (Danecek et al. (2011)) to remove all but the 34 *Anopheles gambiae* and *Anopheles coluzzii* samples. Similarly, we retrieve VCF files and sample IDs for the phase 1 AR3 data release from the 1000 *Anopheles* genome project web site and use VCFtools to remove all but the 150 Burkina Faso samples.

The *Drosophila* samples required more processsing. We retrieve the VCF file for the *Drosophila* Genetics Reference Panel v2 (Huang et al. (2014); Mackay et al. (2012)) from the project web site. We use VCFtools to create a separate VCF file for each chromosome arm (2L, 2R, 3L, 3R, and X). We remove seven samples (lines 348, 350, 358, 385, 392, 395, and 399) that appear to be outliers and then filter each VCF file to only keep biallelic SNPs.

#### Feature Matrix Encoding

Assume that we have *N* samples with *V* positions with biallelic variants. Each position has a reference allele and an alternative allele, and at each position, each sample has one of three genotypes (homozygous reference, homozygous alternate, or heterozygous).

We encode the variants as a feature matrix **X** with dimensions *N* × 3*V*. If sample *i* has the homozygous reference genotype at position *k*, then we set **X**_*i*,3*k*+1_ = 1. If sample *i* has the homozygous alternate genotype at position *k*, then we set **X**_*i*,3*k*+2_ = 1. If sample *i* has the heterozygous genotype at position *k*, then we set **X**_*i*,3*k*+3_ = 1. If the genotype of sample *i* is unknown at position *k*, then we do nothing.

#### Principal Component Analysis (PCA)

Principal component analysis (PCA) of the feature matrix **X** produces a 3*V × P* matrix **W** of principal components and a *N × P* matrix **T** of projected coordinates for the samples such that:

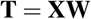

As directly computing PCA would involve computing a 3*V* × 3*V* co-variance matrix, we use a randomized PCA method as implemented in Scikit Learn (Pedregosa et al. (2011)). Whitening is applied to the resulting PCs. We use plots of the explained variance ratios to select relevant PCs.

#### Inferring Karyotypes with K-Means Clustering

Sample karyotypes are inferred by clustering samples using their their projected coordinates (**T**) from PCA. Clustering is performed with the k-means clustering algorithm as implemented in Scikit Learn (Pedregosa et al. (2011)). We choose the number of clusters *K* by clustering the samples with 1-6 clusters, plotting the inertia (or sum-of-squared errors), and visually identifying the “elbow” in the plot. We use the default Scikit Learn settings of 10 runs.

The cluster labels can be represented by a *N × K* matrix **C**. Each sample *i* belongs to one of *K* clusters, indicated by a value of 1 at position **C**_*i*_, _*j*_ where 1 *≤ j ≤ K*.

In cases where we know the karyotypes, we can evaluate the accuracy of the inferred karyotypes from clustering. We generate a confusion matrix for the cluster assignments versus the known karyotypes. From the matrix, we calculate the balanced accuracy of predicting the clusters from the known karyotypes. This set up penalizes situations where the number of clusters is larger than the number of real karyotypes. Balanced accuracy re-weights the accuracy for each class so that each class has equal weight to avoid over-estimating accuracy if poor predictions happen in minority classes.

#### Review of Association Testing

We review associating testing with Logistic Regression models. Likelihood-ratio tests can be used to test for association between variables. The null hypothesis is that knowing the independent variable does not improve the accuracy of predicting the dependent variable, while the alternative hypothesis is that knowing the value of the independent variable does improve accuracy of predictions because the independent variable is associated with the dependent variable.

In our case, we use a Logistic Regression model, which is appropriate when the independent variable is categorical. The equation for a Logistic Regression model is given by:

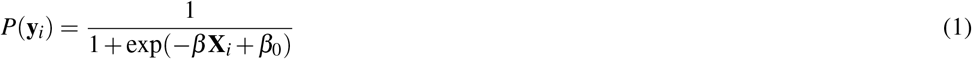

where **y**_*i*_ is value of the dependent variable for sample *i*, **X**_*i*_ is a vector of values for the independent variables for sample *i*, and *β*_0_ is the intercept.

To evaluate the hypothesis, we compare predictions from a pair of models. The alternative model contains the same dependent variables variables as the null model plus the additional independent variable(s) being tested against the dependent variable for association. In our case, the null model only contains an intercept (no independent variables) and the alternative model will contain a single independent variable. In cases where the output variable is categorical rather than binary, a one-versus-all scheme is used. One pair of models is trained for each category and predicts the probability that the value of the independent variable is equal to that category.

After fitting the models, we use the models to predict the independent variable for the samples. From the predictions, we calculate the likelihood for each model. The likelihood for the multinomial Logistic Regression model is given by (Hosmer Jr. et al. (2013)):

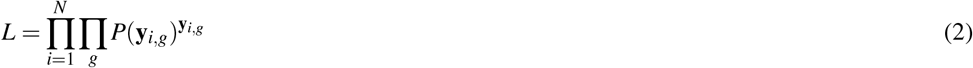

where *g* is the number of categories the dependent variable can take on.

To perform the likelihood-ratio test, the difference *G* between the log likelihoods of the two sets of models is calculated by:

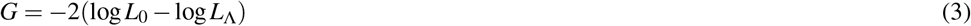

where *L*_0_ and *L*_Λ_ are the likelihoods of the null and alternative models, respectively. The *p*-value for the difference in log likelihoods is calculated using the *χ*2 distribution:

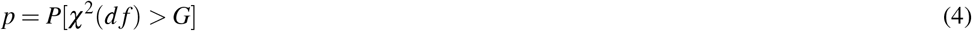

where *d f* is the difference in the number of degrees of freedom (weights) between the two models. Scikit Learn is used; we train the models using Stochastic Gradient Descent (SGD) for 10,000 epochs, the log likelihood, *L*_2_ regularization using the `SGDClassifier` class. All other parameters are left at their defaults. The log likelihoods are calculated with the `log_loss` function (normalize set to `False`). We implement functionality for calculating *G* and estimating the *p*-value using Scipy.

#### Localizing Inversions with Cluster-SNP Association Tests

After karyotypes are inferred with clustering, we perform association tests between each SNP and the samples’ cluster labels. The cluster labels are used as the independent variables (**y**), while the genotypes of the SNPs are used as the independent variables (**X**).

It is common for genotypes in insect SNP data to be unknown (uncalled). We use our approach from Nowling and Emrich (2018a,b) to adjust the association tests to avoid bias. For fitting the models, we deterministically up-sample the samples (one copy for each possible genotype). In particular, if we have *M* genotypes, we create *M* copies of each sample. (In our case, *M* = 3 since we are working with biallelic SNPs with three genotypes.) If the genotype is known, the copies have the same genotype as the original. Otherwise, we make the conservative assumption that there is an uninformative (uniform) prior over the genotypes and impute the copies so that there is a one-to-one relationship between the copies and possible genotypes. Additionally, we fix the intercept to the class probabilities and did not allow it to be changed during fitting. For prediction and evaluation of the likelihood, we use original input data.

#### Localizing Inversions with PC-SNP Association Tests

In Nowling and Emrich (2018c), we described a second approach for localizing inversions in which association tests are performed between each SNP and the samples’ PC projected coordinates (*T*) from PCA. A single association test is performed for each combination of principal component (PC) *j* and SNP position *k*, using the coordinate *T*_*i*_, _*j*_ for sample *i* along PC *j* as the independent variable. As the SNPs are encoded as categorical variables, three dependent variables (one for each genotype) are used for each SNP. We employ three Logistic Regression models, one for each genotype, in a one-versus-all scheme.

As the SNPs are the dependent variables, we need a different strategy for handling missing genotypes. We review the method we proposed in Nowling and Emrich (2018c). We deterministically up-sample the samples (one copy for each genotype). In particular, if we have *M* genotypes, we create *M* copies of each sample. (In our case, *M* = 3 since we are working with biallelic SNPs with three genotypes.) If the genotype is known, the copies have the same genotype as the original. Otherwise, we make the conservative assumption that there is an uninformative (uniform) prior over the genotypes and impute the copies so that there is a one-to-one relationship between the copies and possible genotypes. We also fix the intercept to the class probabilities and did not allow it to change during fitting. Note that unlike the approach for the cluster-SNP association tests, the up-sampled data are used for both fitting the models and in predictions for the calculations of the likelihoods.

Since we increased the number of samples, we need to weight the samples so that the calculated *p*-values are consistent with the original number of samples. The modified likelihood function is then:

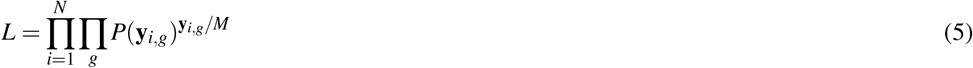

#### Software Implementation

We implement our method in Asaph, our open-source toolkit for variant analysis. Asaph is implemented in Python using Numpy / Scipy (Walt et al. (2011)), Matlotlib (Hunter (2007)), and Scikit-Learn (Pedregosa et al. (2011)) and is available at https://github.com/rnowling/asaph under the Apache Public License v2.

## RESULTS

### Analysis of Simulated Inversions

We first simulate 500 diploid individuals with a single 2 Mb chromosome containing a single inversion spanning 0.75Mb to 1.25Mb using invertFREGENE (O’Reilly et al. (2010)). The inverted and standard homozygotes each corresponded to 25% of the samples, while 50% of the samples are heterozygous.

Explained variance ratios for the PCA of the invertFREGENE data indicates that three PCs are needed to explain most of the variation, but cluster structure was only present in the projection plot for PCs 1 and 2 (see Figure 1a-c). K-means identifies three clusters (see Figure 1d). The balanced accuracy for predicting clusters assignments from karyotype labels was 100.0%, which indicates a perfect one-to-one relationship between the three clusters and three inversion karyotypes. Significantly, a Manhattan plot of the SNPs’ associations with the cluster labels indicate the presence of the inversion in the expected location (see Figure 1e).

**Figure 1.**
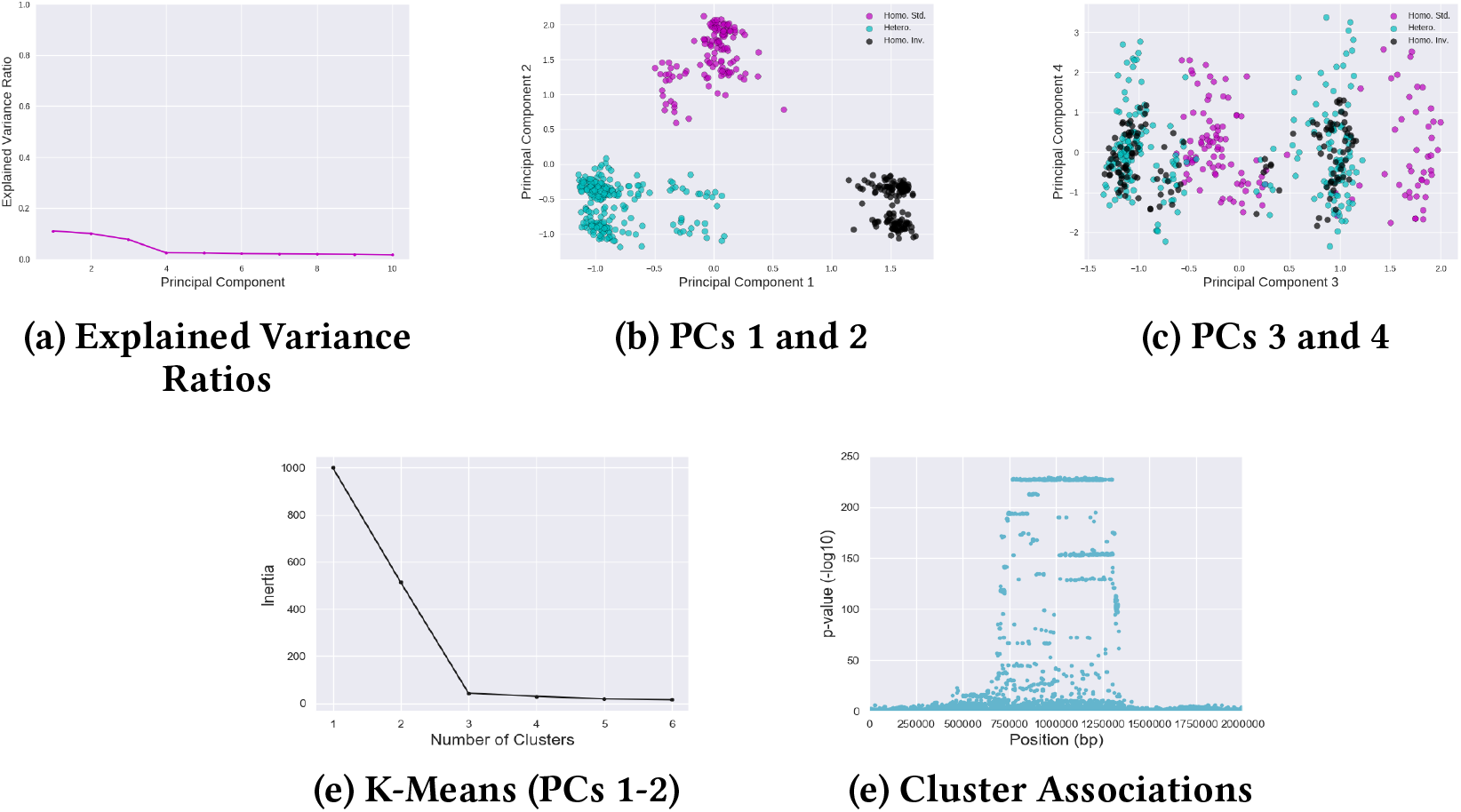
Analysis of SNPs from 500 Individuals Simulated with invertFREGENE with PCA, Clustering, and Cluster-SNP Association Tests. (a) Explained variance ratios, (b–c) PCA projection plots, and (d–f) Manhattan plots from Cluster-SNP association tests.

These simulations confirmed that PCA and k-means clustering of SNPs can be used to infer inversion karyotypes by validating the assigned clusters against the known karyotype labels. Further, association tests between the clusters and SNPs can localize the inversion along the chromosome.

### Analysis of *Drosophila* Inversions

Samples in the *Drosophila* Genetics Reference Panel 2 (DGRP2) data contain multiple inversion karyotypes and are drawn from a single population. Only five inversions are present in five or more samples (Huang et al. (2014)). The 2L and 2R chromosome arms each contain a single inversion (*ln(2L)t*, *ln(2R)NS*) and all three orientations are present for each inversion. Three inversions (*ln(3R)P*, *ln(3R)K*, and *ln(3R)Mo*) are present on the 3R chromosome arm. The three inversions overlap and the inverted orientations are nearly mutually exclusive in the DGRP2 samples (see Tables 1–3).

**Table 1.** Co-occurrences of *In(3R)Mo* and *In(3R)K* Inversion Karyotypes in 198 *Drosophila* Samples

|  |  | *In(3R)Mo* |  |  |
| --- | --- | --- | --- | --- |
|  |  | Homo. Std. | Hetero. | Homo. Inv |
| *In(3R)K* | Homo. Std. | 167 | 8 | 17 |
|  | Hetero. | 9 | 1 | 0 |
|  | Homo. Inv | 3 | 0 | 0 |

**Table 2.** Co-occurrences of *In(3R)Mo* and *In(3R)P* Inversion Karyotypes in 198 *Drosophila* Samples

|  |  | *In(3R)Mo* |  |  |
| --- | --- | --- | --- | --- |
|  |  | Homo. Std. | Hetero. | Homo. Inv |
| *In(3R)P* | Homo. Std. | 169 | 9 | 17 |
|  | Hetero. | 6 | 0 | 0 |
|  | Homo. Inv | 4 | 0 | 0 |

**Table 3.** Co-occurrences of *In(3R)K* and *In(3R)P* Inversion Karyotypes in 198 *Drosophila* Samples

|  |  | *In(3R)K* |  |  |
| --- | --- | --- | --- | --- |
|  |  | Homo. Std. | Hetero. | Homo. Inv |
| *In(3R)P* | Homo. Std. | 182 | 10 | 3 |
|  | Hetero. | 6 | 0 | 0 |
|  | Homo. Inv | 4 | 0 | 0 |

**Table 4.** Association Tests Between Principal Components and Inversion Karyotypes of *Drosophila* Samples. PCA was performed separately for each chromosome, so the PC columns refer to the PCs for the chromosome of the given inversion.

|  | Comparison | PC1 | PC2 |
| --- | --- | --- | --- |
| **In(2L)t** | Inverted vs Not | x |  |
| **In(2L)t** | Homo. Inverted vs Rest |  |  |
| **In(2L)t** | Hetero. vs Rest |  | x |
| **In(2R)ns** | Inverted vs Not | x |  |
| **In(2R)ns** | Homo. Inverted vs Rest |  |  |
| **In(2R)ns** | Hetero. vs Rest |  | x |
| **In(3R)P** | Inverted vs Not |  |  |
| **In(3R)P** | Homo. Inverted vs Rest |  |  |
| **In(3R)P** | Hetero. vs Rest |  |  |
| **In(3R)K** | Inverted vs Not |  |  |
| **In(3R)K** | Homo. Inverted vs Rest |  |  |
| **In(3R)K** | Hetero. vs Rest | x | x |
| **In(3R)Mo** | Inverted vs Not |  |  |
| **In(3R)Mo** | Homo. Inverted vs Rest | x |  |
| **In(3R)Mo** | Hetero. vs Rest |  |  |

The explained variance ratios from PCAs of the *Drosophila* 2L and 2R SNPs indicates that two PCs per arm are needed to explain most of the variation. In each case, k-means identifies three clusters. The Manhattan plots of the SNPs’ associations with the cluster labels indicates that the clusters are capturing the inversions (see Figures 2d and 3d). The clusters are strongly associated with the karyotypes labels; balanced accuracies for predicting the cluster assignments from the karyotype labels are 93.3% (*In(2L)t*) and 94.4% (*In(2R)NS*), respectively.

**Figure 2.**
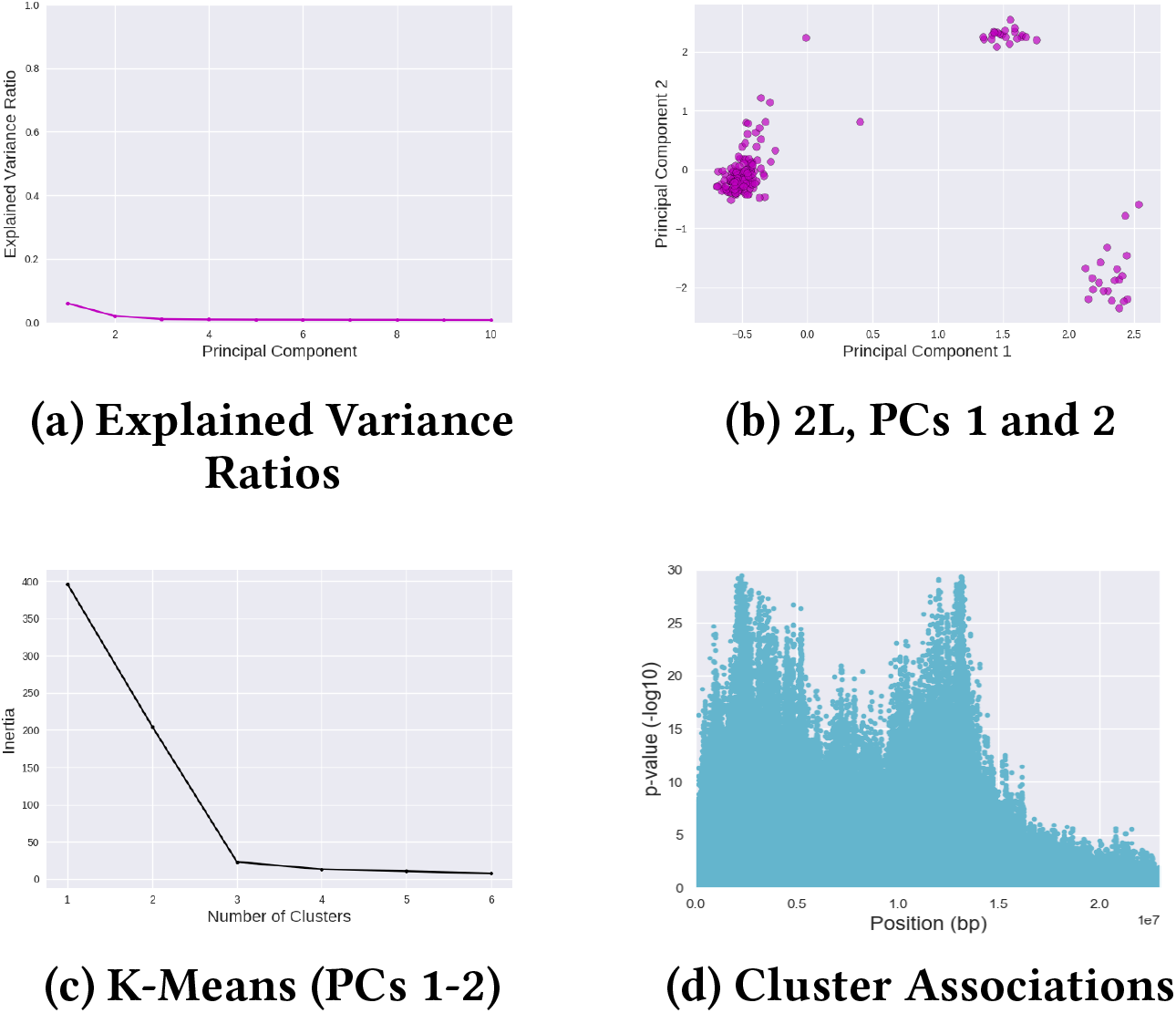
Analysis of 2L Chromosome Arm SNPs of 198*Drosophila* Samples with PCA, Clustering, and Cluster-SNP Association Tests. (a) Explained variance ratios, (b) PCA projection plot, (c) Inertia plot for K-Means clustering, and (d) Manhattan plots from Cluster-SNP association tests.

**Figure 3.**
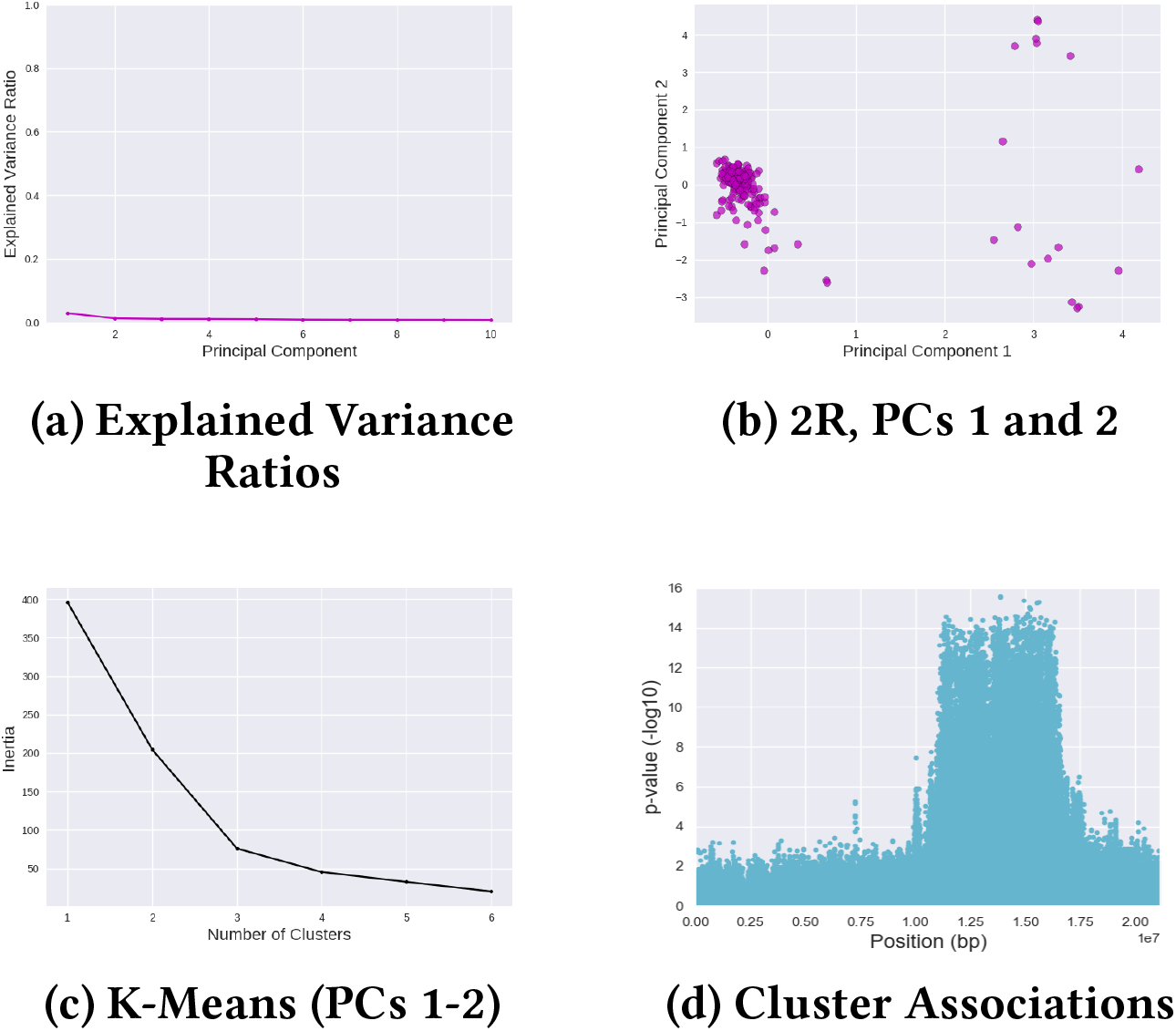
Analysis of 2R Chromosome Arm SNPs of 198*Drosophila* Samples with PCA, Clustering, and Cluster-SNP Association Tests. (a) Explained variance ratios, (b) PCA projection plot, (c) Inertia plot for K-Means clustering, and (d) Manhattan plots from Cluster-SNP association tests.

The inversion story for the 3R chromosome arm is more complicated. Three inversions (*In(3R)P*, *In(3R)K*, and *In(3R)Mo*) on 3R are present in more than five of the DGRP2 samples (Huang et al. (2014)), and although these three inversions overlap the inverted orientations are nearly mutually exclusive in the DGRP2 samples (see Tables 1–3). For these data PCA and clustering are not able to accurately karyotype; two PCs explained most of the variation (see Figure 4a) and k-means clustering using PCs 1 and 2 finds three clusters (see Figure 4c), but the clusters do not correlate with the orientations of any single inversion. Balanced accuracies for predicting clusters assignments from karyotype labels are 55.0% (*In(3R)K*), 60.7% (*In(3R)mo*), and 43.3% (*In(3R)p*).

**Figure 4.**
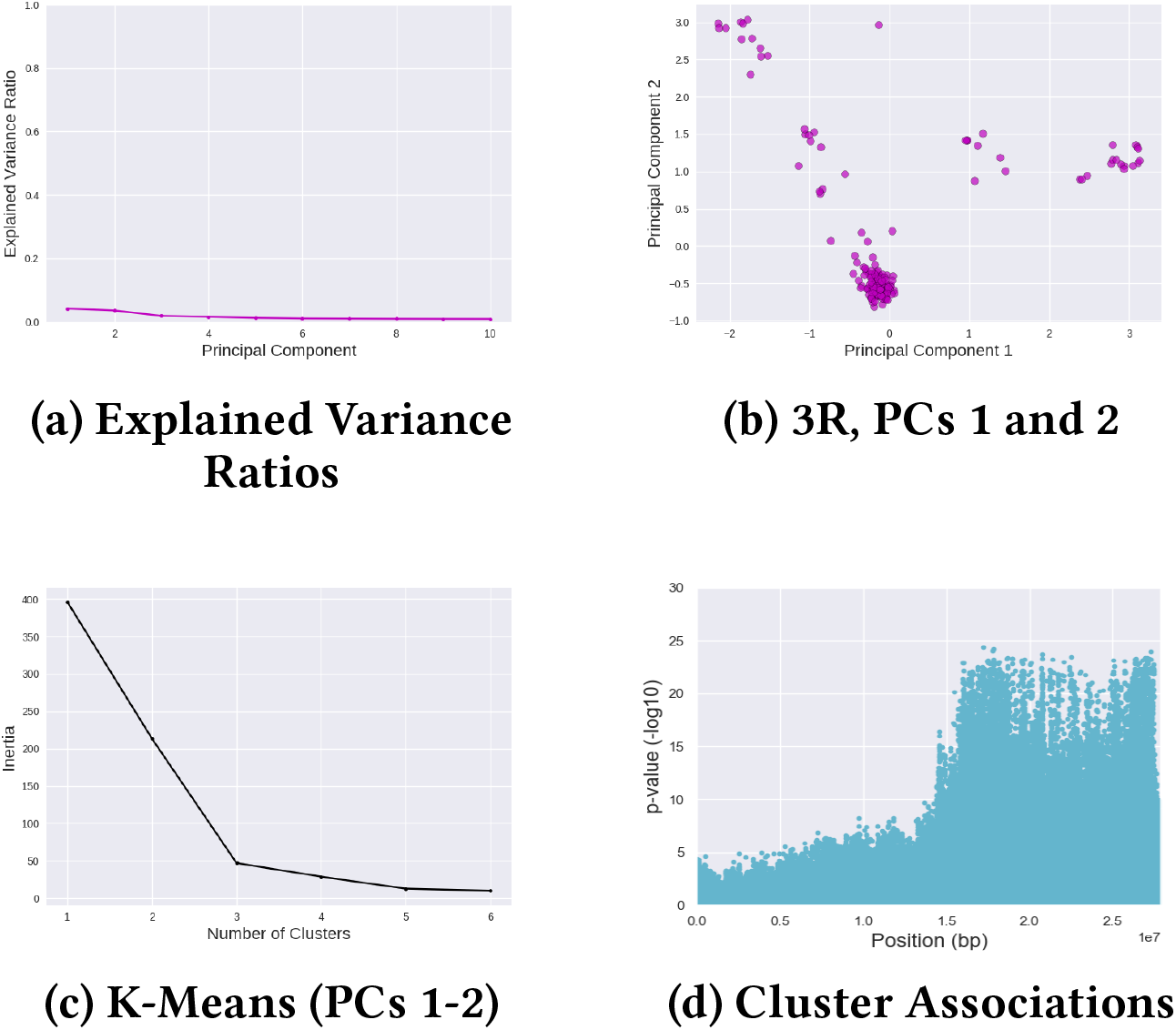
Analysis of 3R Chromosome Arm SNPs of 198*Drosophila* Samples with PCA, Clustering, and Cluster-SNP Association Tests. (a) Explained variance ratios, (b) PCA projection plot, (c) Inertia plot for K-Means clustering, and (d) Manhattan plots from Cluster-SNP association tests.

SNP-cluster association tests, however, are able to localize the region on 3R containing the *In(3R)K* and *In(3R)Mo* inversions but are unable to disambiguate the overlapping inversions. In the Manhattan plots, SNPs associated with the clusters are localized to a large region starting at 15 Mbp and span the rest of the arm, and as such the region appears as to contain one large inversion (see Figure 4d).

Association tests between the PCs and karyotype labels offer an explanation. The first PC divides the two highest-frequency orientations, while the second PC divides the third highest-frequency orientation from the the rest. With multiple mutually-exclusive inversions, however, the two highest-frequency, mutually-exclusive orientations (homozygous inverted *In(3R)Mo* and heterozygous *In(3R)K*) do not belong to the same inversion. Hence, 3R-PC 1 divides the samples with the homozygous inverted orientation of *In(3R)Mo* and heterozygous inversion of *In(3R)K* from the rest. As a result, PCA methods are not successful on 3R because the results could be interpreted computationally as a single inversion when given these three mutually-exclusive but overlapping inversions.

### Analysis of inversions found in less closely related samples

We also analyze Burkina Faso *Anopheles gambiae* and *Anopheles coluzzii* samples from the 1000 *Anopheles* genomes project. The samples samples were karyotyped for the 2La and 2Rb inversions. Not all karyotypes are present for the 2La inversion, however, which complicates detection and karyotyping because none of the samples are homozygous for the standard 2La karyotype and only a single *An. coluzzii* sample is heterozygous (see Table 5).

**Table 5.** Occurrences of 2La Inversion Karyotypes By Species for 150 Burkina Faso *Anopheles* Samples

|  | 2La |  |  |
| --- | --- | --- | --- |
|  | Homo. Std. | Hetero. | Homo. Inv |
| *An. coluzzii* | 0 | 1 | 68 |
| *An. gambiae* | 0 | 15 | 66 |

**Table 6.** Occurrences of 2Rb Inversion Karyotypes By Species for 150 Burkina Faso *Anopheles* Samples

|  | 2Rb |  |  |
| --- | --- | --- | --- |
|  | Homo. Std. | Hetero. | Homo. Inv |
| *An. coluzzii* | 33 | 29 | 7 |
| *An. gambiae* | 2 | 24 | 55 |

We repeat the approach of inferring karyotypes to the 2L and 2R chromosome arms of a total of 150 Burkina Faso *Anopheles gambiae* and *Anopheles coluzzii* samples. PCA of the samples detects differences between species and inversion karyotypes as previously reported (see Figures 6a and 5a). Because the resulting clusters combine species and karyotype, isolation of the inversion effects and localization of the inversions is difficult using this method.

We therefore divide the samples by species and perform PCA on each species separately. Since only a single *An. coluzzii* sample is inverted for 2La, none of the PCs had large explained variance ratios and we are unable to use PCA to karyotype these *An. coluzzii* samples or localize the 2La inversion. For *An. gambiae*, k-means identifies two clusters, corresponding to the homozygous inverted and heterozygous orientations (balanced accuracy of 100.0%). The location of the 2La inversion is clearly indicated based on a Manhattan plot generated from the association test results (see Figure 5f).

**Figure 5.**
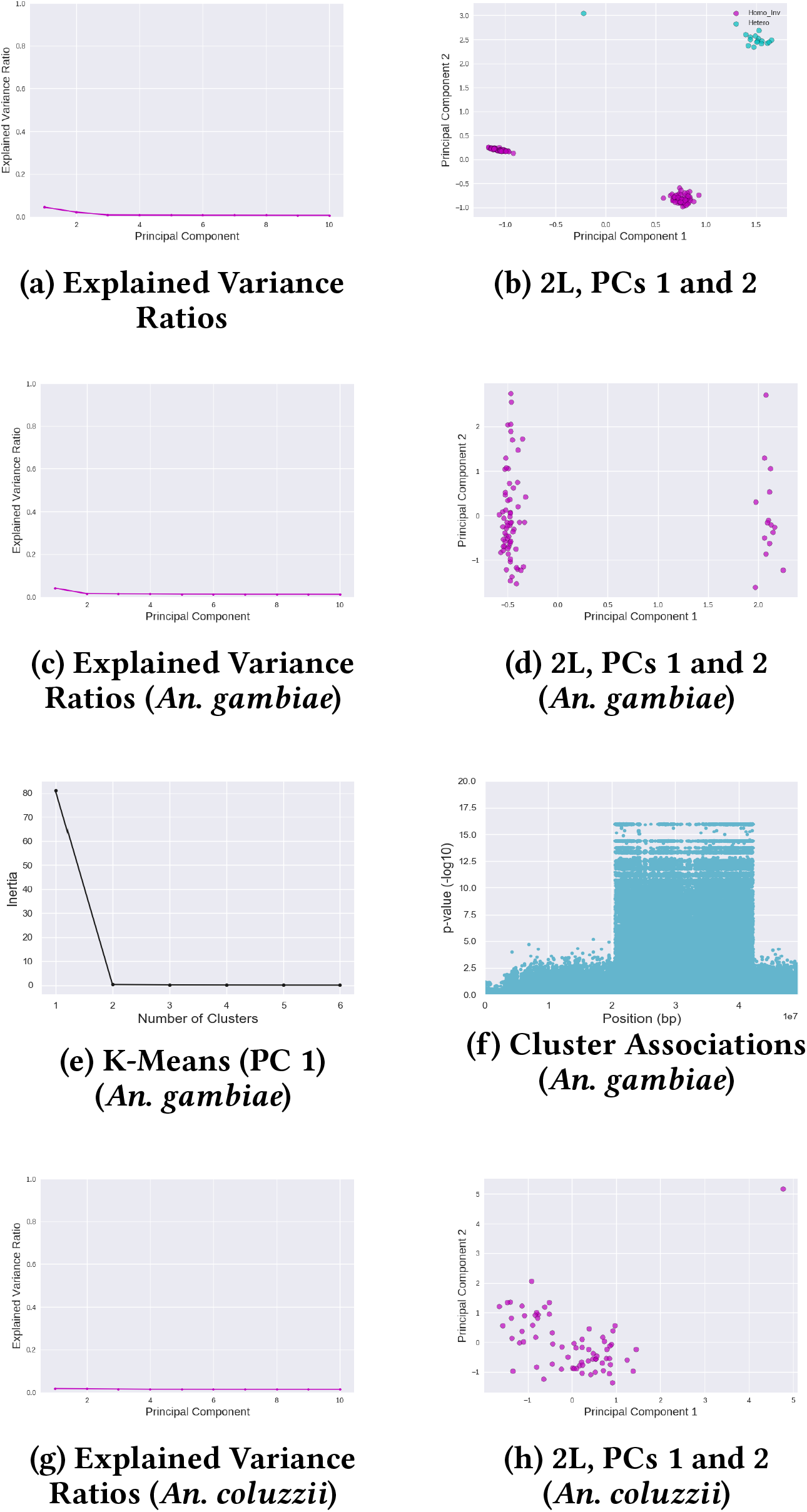
Analysis of 2L Chromosome Arm of 150 Burkina Faso Anopheles Samples with PCA, Clustering, and Cluster-SNP Association Tests. The samples clustered by species and karyotype, so samples were divided and re-analyzed by species. (a) Explained variance ratios for all samples, (b) PCA projection plot for all samples, (c) explained variance ratios for *An. gambiae* samples, (d) PCA projection plot for *An. gambiae* samples, (e) Inertia plot for K-Means clustering of *An. gambiae* samples, (f) Manhattan plots from Cluster-SNP association tests for *An. gambiae* samples, (g) explained variance ratios for *An. coluzzii* samples, and (h) PCA projection plot for *An. coluzzii* samples.

**Figure 6.**
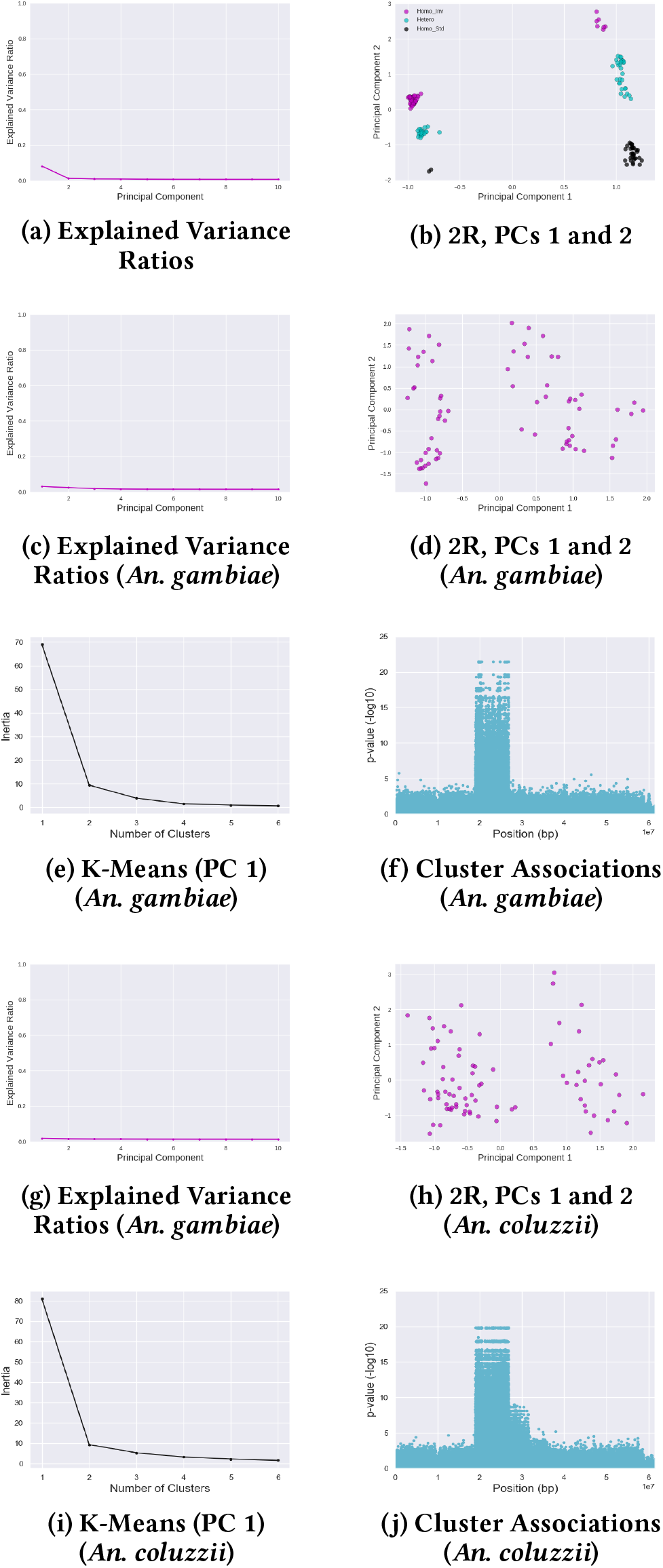
Analysis of 2R Chromosome Arm of 150 Burkina Faso *Anopheles* Samples with PCA, Clustering, and Cluster-SNP Association Tests. The samples clustered by species and karyotype, so samples were divided and re-analyzed by species. (a) Explained variance ratios for all samples, (b) PCA projection plot for all samples, (c) explained variance ratios for *An. gambiae* samples, (d) PCA projection plot for *An. gambiae* samples, (e) Inertia plot for K-Means clustering of *An. gambiae* samples, (f) Manhattan plots from Cluster-SNP association tests for *An. gambiae* samples, (g) explained variance ratios for *An. coluzzii* samples, (h) PCA projection plot for *An. coluzzii* samples, (i) Inertia plot for K-Means clustering of *An. coluzzii* samples, and (j) Manhattan plots from Cluster-SNP association tests for *An. coluzzii* samples.

For 2R, two PCs explains most of the variance for the *An. coluzzii* samples, while one PC explains most of the variance for the *An. gambiae* samples; in both cases, we find that using only the first PC produces the best clustering results. K-means identifies two clusters of *An. gambiae* samples, which correlate perfectly with the homozygous inverted and heterozygous orientations, and the balanced accuracy for predicting clusters assignments from karyotype labels is also 100.0% for *An. gambiae* and *An. coluzzii* even though the two homozygous standard samples are not detected as a separate cluster. Manhattan plots generated from the SNP-cluster association results successfully localizes the 2Rb inversion in both species (see Figures 6f and 6j).

Notably, the Manhattan plots suggest that the 2Rc inversion (Main et al. (2015)) may also be present in some of the *An. coluzzii* samples even though they were not karyotyped for 2R inversions other than 2Rb. When the 2Rb and 2Rc inversions appear together, they are designated as the 2Rbc system (Caputo et al. (2014)). The presence of 2Rc (2Rbc) in some of the *An. coluzzii* samples may explain why the karyotypes from the two species did not cluster together along PC 2 when the 150 samples are analyzed together.

### Multiple Inversions, Multiple Species, Multiple Populations

We apply our approach to the analysis of 34 *Anopheles gambiae* and *Anopheles coluzzii* samples from four geographic locations (Burkina Faso, Cameroon, Mali, and Tanzania) (Fontaine et al. (2015)). These samples were karyotyped for the 2La inversion, but not inversions on the 2R chromosome arm.

The 2La karyotype labels between the 34 *Anopheles* and 150 Burkina Faso *Anopheles* samples may not be consistent: 2La homozygous inverted orientation is not observed among the 7 Burkina Faso samples from the 34 total *Anopheles* samples, while the 2La homozygous standard orientation is not observed among the 150 Burkina Faso *Anopheles* samples (see Tables 5 and 8).

The 2La inversion forms are associated with both species and locations. Samples from Cameroon are primarily homozygous for the inverted orientation, while samples from Burkina Faso and Mali are primarily homozygous for the standard orientation (see Table 7). Five samples from across locations are heterozygous. All three orientations were observed in *An. gambiae* samples, while *An. coluzzii* samples are homozygous for either the standard or inverted orientations (see Table 8). Due in part to the small sample size, we conclude that the inversion karyotypes are not easily separated from the species or geographic location in this initial analysis.

**Table 7.** Occurrences of 2La Inversion Karyotypes By Location for 34 *Anopheles* Samples

|  | 2La |  |  |
| --- | --- | --- | --- |
|  | Homo. Std. | Hetero. | Homo. Inv |
| Burkina Faso | 5 | 2 | 0 |
| Cameroon | 0 | 1 | 15 |
| Mali | 8 | 0 | 0 |
| Tanzania | 0 | 2 | 1 |

**Table 8.** Occurrences of 2La Inversion Karyotypes By Species for 34 *Anopheles* Samples

|  | 2La |  |  |
| --- | --- | --- | --- |
|  | Homo. Std. | Hetero. | Homo. Inv |
| *An. coluzzii* | 3 | 0 | 8 |
| *An. gambiae* | 10 | 5 | 8 |

Two PCs explain most of the variance for the 2L SNPs. Using PC 1, k-means is able to identify three clusters. The balanced accuracy for predicting clusters assignments from karyotype labels is 100.0%. Manhattan plots from the SNP-cluster association tests successfully localizes the 2La inversion (see Figure 7).

**Figure 7.**
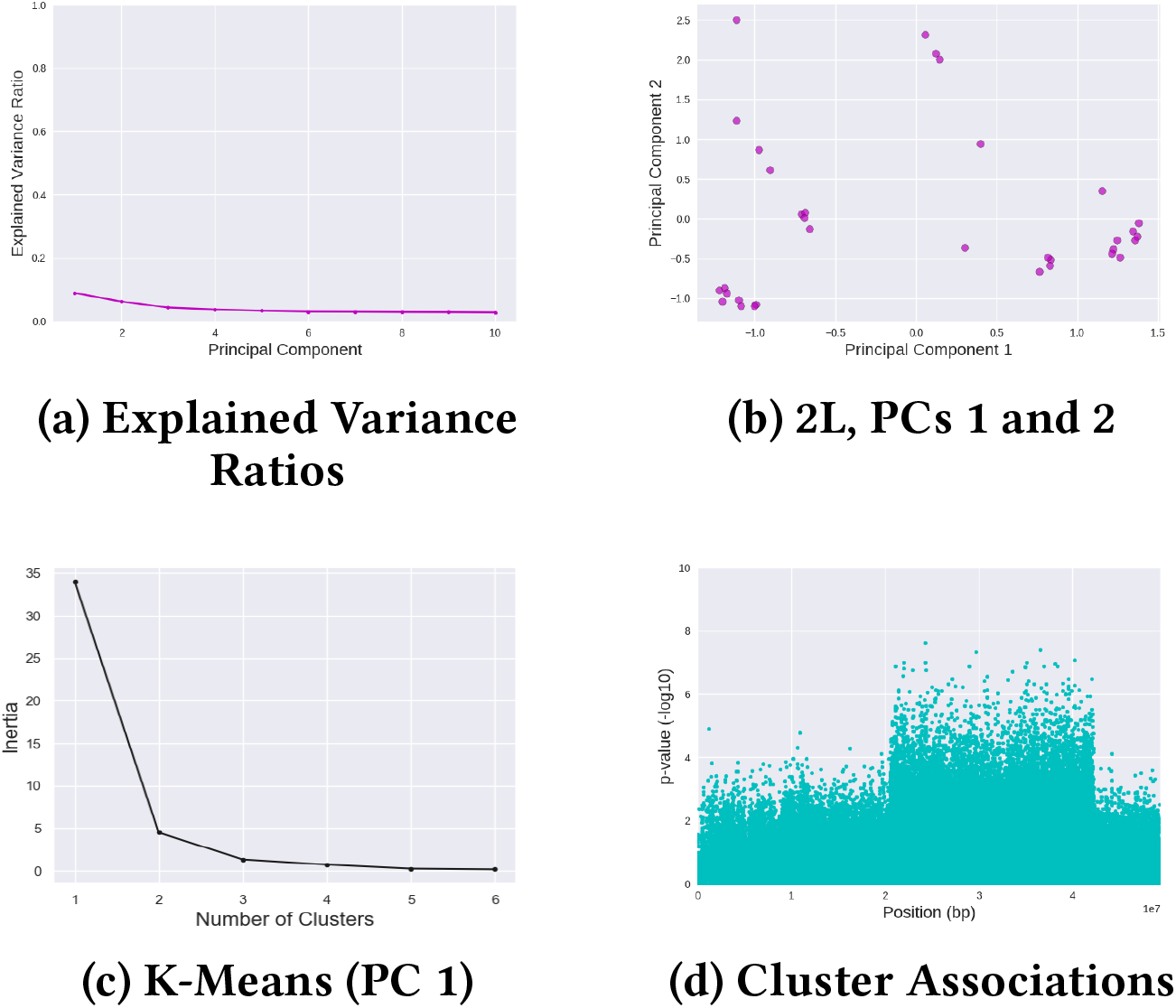
Analysis of 2L Chromosome Arm of 34 *Anopheles* Samples with PCA, Clustering, and Cluster-SNP Association Tests. (a) Explained variance ratios, (b) PCA projection plot, (c) Inertia plot for K-Means clustering, and (d) Manhattan plots from Cluster-SNP association tests.

We also identify inversions on 2R (see Figure 8). Four PCs explain most of the variance. K-mean identifies three clusters using PCs 1 and 2. Association tests with the clusters labels from PCs 1 and 2 identify potential inversions. There are multiple inversions (e.g., 2Rj, 2Rb, 2Rc, and 2Rj) on 2R, including several (e.g., 2Rbk, 2Rcu, 2Rbu, and 2Rd) that overlap (Main et al. (2015); Caputo et al. (2014)). The Manhattan plot shows associated SNPs in the 2Rj inversion region near the front of the chromosome arm. The second set of associated SNPs do not overlap entirely with the 2Rb inversion and could potentially belong to the 2Rbk or 2Rcu inversion systems (Caputo et al. (2014)). The eight *An. gambiae* samples from Mali formed one of the three clusters, suggesting that the potential inversions captured are present (or absent) predominantly in Mali.

**Figure 8.**
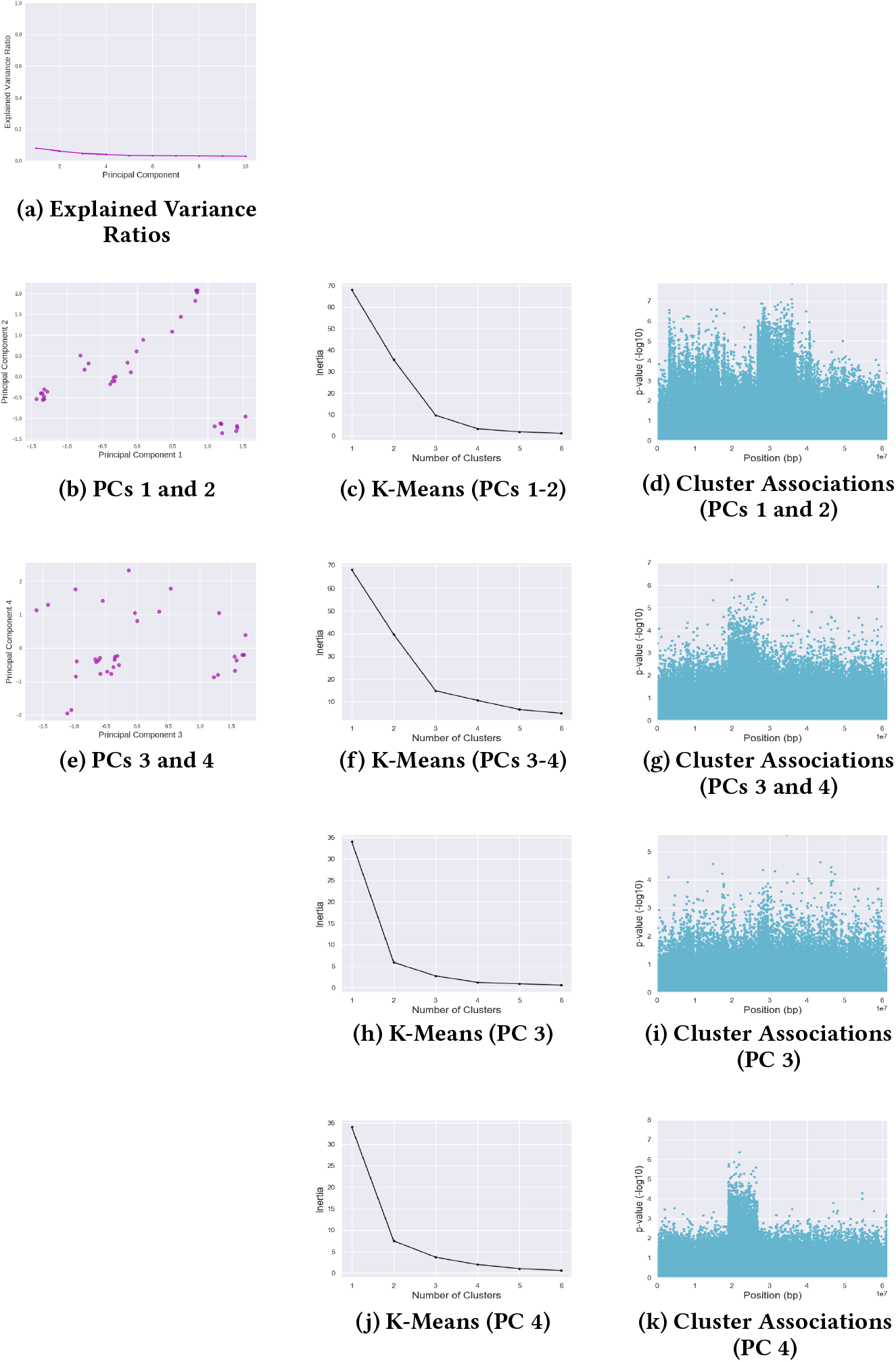
Analysis of 2R Chromosome Arm of 34 *Anopheles* Samples with PCA, Clustering, and Cluster-SNP Association Tests. The explained variance analysis indicates that first 3-4 PCs were significant, so PCs 1 and 2 were analyzed followed by PCs 3 and 4. (a) Explained variance ratios, (b) PCA projection plot for PCs 1-2, (c) Inertia plot for K-Means clustering (PCs 1-2), (d) Manhattan plots from Cluster-SNP association tests for PCs 1-2, (e) PCA projection plot for PC 3 and 4, (f) Inertia plot for K-Means clustering (PCs 3-4), (g) Manhattan plots from Cluster-SNP association tests for PCs 3-4, (h) Inertia plot for K-Means clustering (PC 3), (i) Manhattan plots from Cluster-SNP association tests for PC 3, (j) Inertia plot for K-Means clustering (PC 4), and (k) Manhattan plots from Cluster-SNP association tests for PC 4.

Three clusters are identified using PCs 3 and 4. The 2Rb inversion is present in the corresponding Manhattan plot, although not clearly. We re-clustered the samples separately for each PC. Two to three clusters are identified for each PC. The Manhattan plot for the PC 4 clusters reveals the 2Rb inversion clearly, while the Manhattan plot for the PC 3 clusters does not indicate an inversion. PC 4 captures the 2Rb inversion, while PC 3 likely captures something other than an inversion. Although these samples are not karyotyped for 2R inversions, the presence of the 2Rb inversion is expected based on its presence in the larger 150 Burkina Faso set of samples.

### Comparison to PC-SNP Association Tests

In previous sections, we evaluate PCA and clustering for inferring inversion karyotypes and association tests with the cluster labels for localizing inversions. We previously described an alternative approach in which association tests are performed directly against the projected PC coordinates (no intermediate clustering step) (Nowling and Emrich (2018c)). PC-SNP association tests are able to detect and localize inversions but unable to infer karyotypes. For completeness we re-analyze the above data using our alternative PC-SNP association test approach.

For the cases with a single inversion and no population structure, the two methods are equal in their ability to localize inversions. The inversion in the invertFREGENE simulation is localized by PCs 1 and 2 (see Figures 9a and 9b); PC 3 captures an unrelated effect. The *Drosophila In(2L)t* and *In(2R)NS* inversions are localized by the first PC for each chromosome arm (see Figures 10a and 9c); the second PCs capture differences between homozygous and heterozygous karyotypes (see Table 4), but do not localize the inversion.

**Figure 9.**
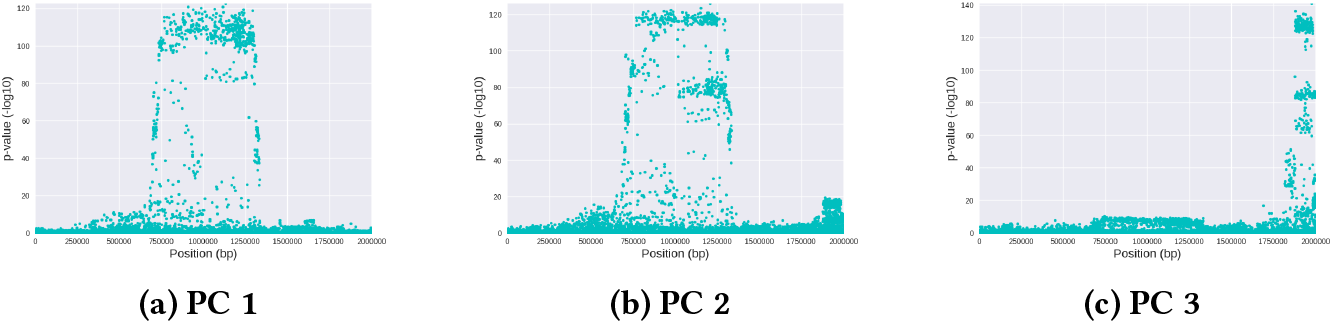
Manhattan Plots from PC-SNP Associations for invertFREGENE Samples.

**Figure 10.**
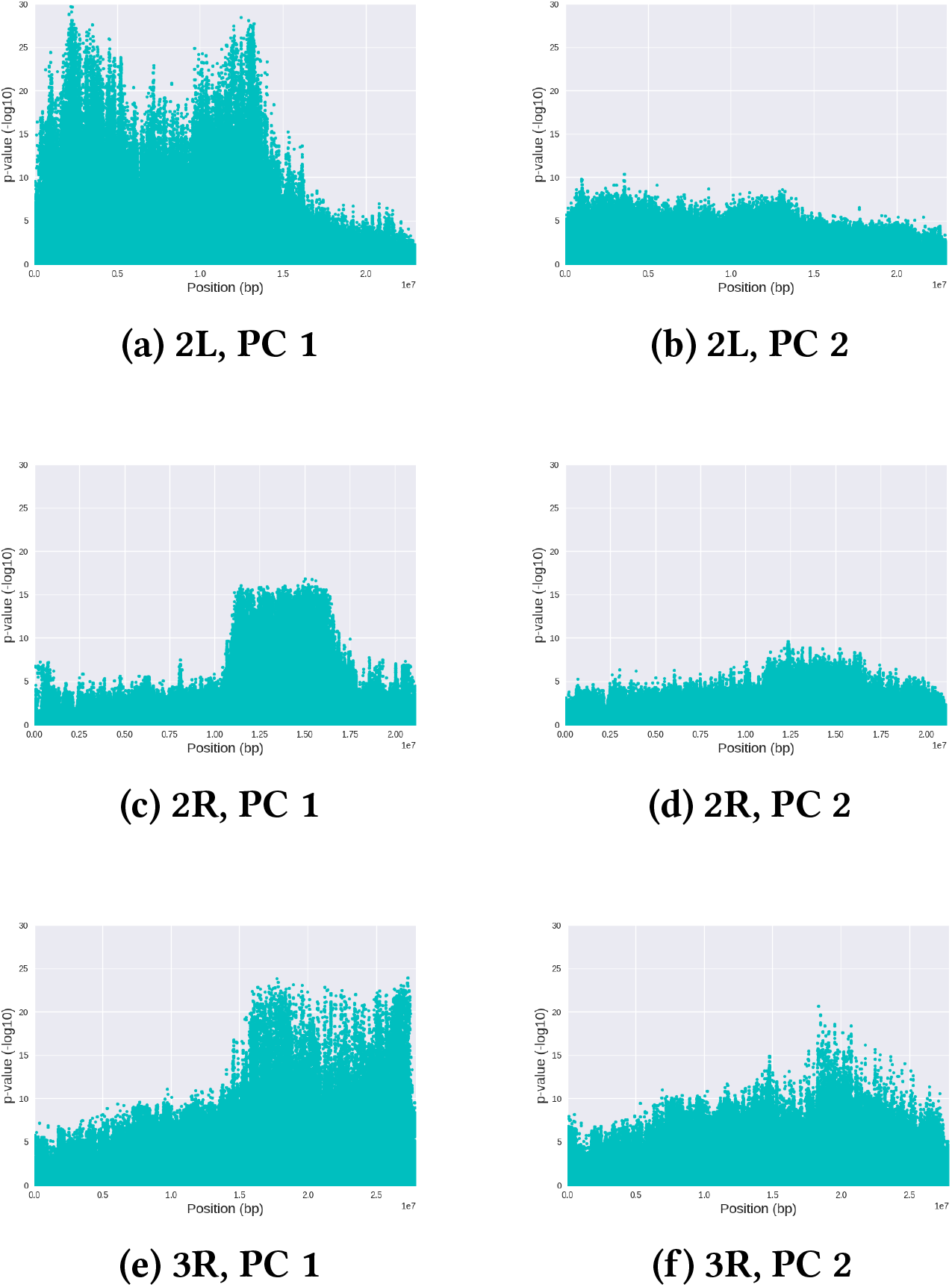
Manhattan Plots from PC-SNP Associations for 198 *Drosophila* Samples.

PC-SNP association tests are more robust to population structure and confounding factors. For the 150 Burkina Faso samples, we observed that the PC 1 captures differences between species, while PC 2 captures the inversions. Accordingly, association tests against the second PCs localize the 2La and 2Rb inversions (see Figures 11b and 11d). For the 34 *Anopheles* samples, the 2La inversion is localized by association tests against 2L-PC 1 (see Figure 11a), the 2Rb inversion is localized by 2R-PC 4 (see Figure 11h), and as hypothesized earlier, 2R-PC 2 is capturing inversions what might be the 2Ru and 2Rcu or 2Rbk inversion systems (see Figure 11d).

**Figure 11.**
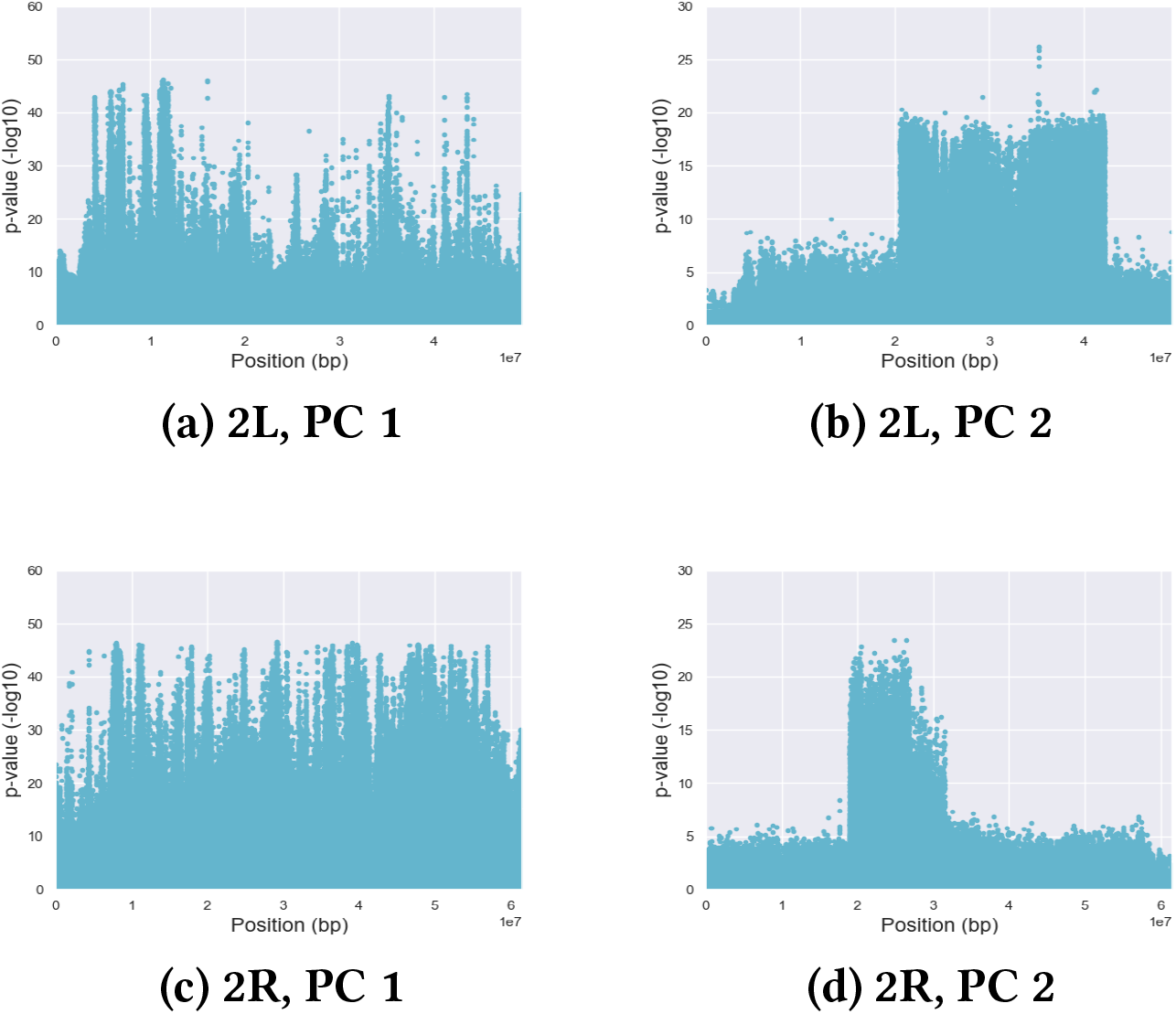
Manhattan Plots from PC-SNP Associations for 150 Burkina Faso *Anopheles* Samples.

**Figure 12.**
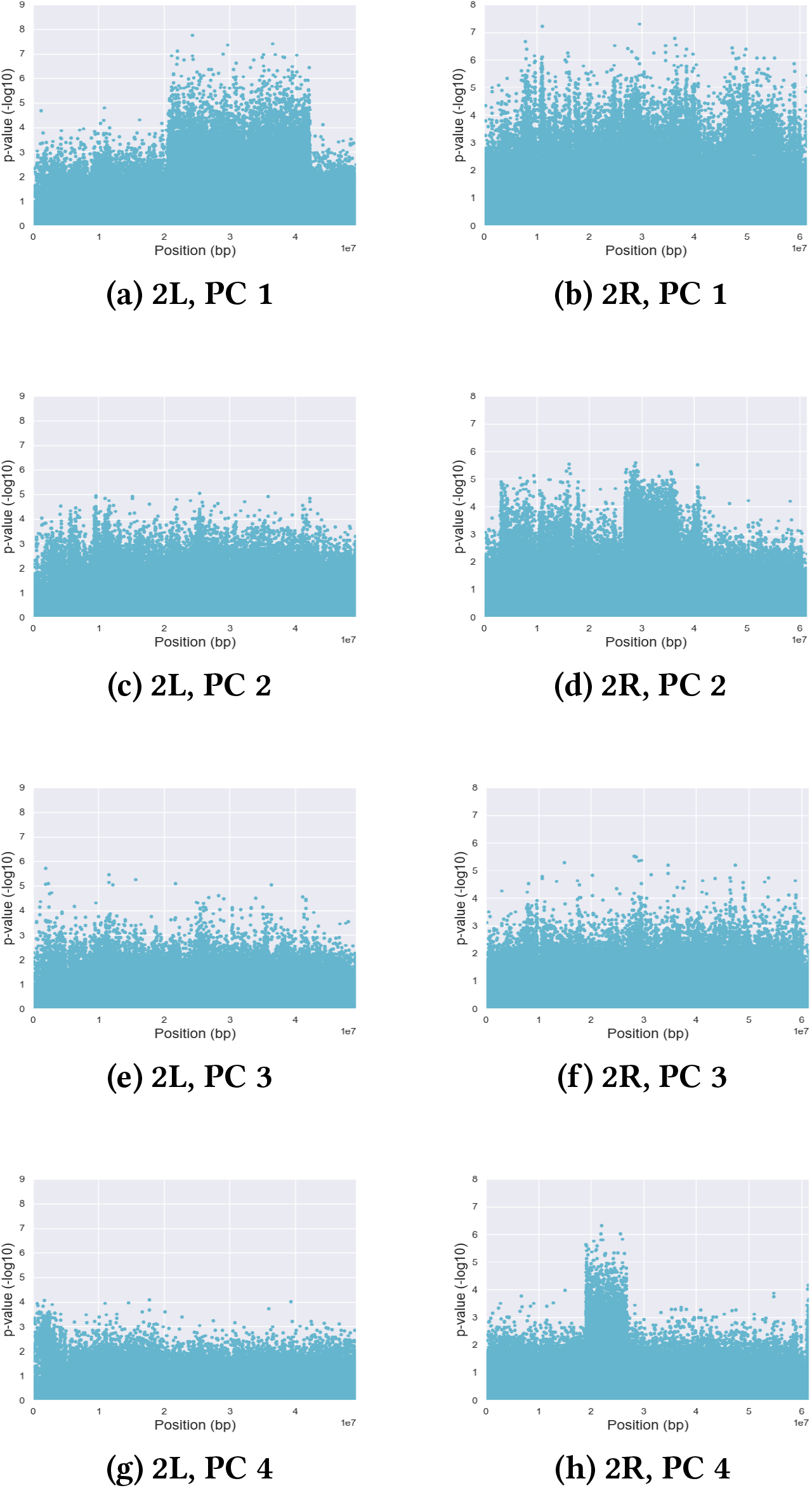
Manhattan Plots from PC-SNP Associations for 34*Anopheles* **Samples**.

Finally, we observe that the association tests against the projected coordinates do not resolve the ambiguity from the multiple overlapping inversions on the 3R chromosome arm of the 198 *Drosophila* samples. Only 3R-PC 1 appears to localize an inversion (see Figure 10e), and the enriched region appears as a single inversion.

## DISCUSSION

We evaluate PCA-based frameworks for detecting, localizing, and karyotyping inversions from SNPs. Although both approaches (cluster-SNP associations and PC-SNP associations) are practical and useful for identifying large inversions, there are trade offs. While the cluster-based approach is able to infer karyotypes, it requires choosing an appropriate combination of PCs and the right number of clusters. The second approach has fewer requirements but cannot infer karyotypes.

When applied to simulated and real data (*Drosophila* 2L and 2R chromosome arms) with a single inversion and a single population, both methods readily detect and localize the inversions while the cluster-based approaches are able to correctly infer karyotypes.

Sample data with more complicated inversions and population structure proved more challenging. While the *Drosophila* 3R chromosome arm has three overlapping and mutually-exclusive inversions, PCA only indicates one inversion with three karyotypes. Without prior knowledge of the karyotypes, the results from PCA could be misinterpreted. Using data with multiple, closely-related species, PCA analysis detects the differences in species as well as the inversions. We found it necessary to analyze the 150 Burkina Faso *Anopheles* samples separately by species to accurately resolve the karyotypes and inversions. We observe the expected 2Rb inversion, but we also observe the presence of the 2Rc inversion within some *An. coluzzii* samples. We note that not knowing *a priori* that the 2Rc inversion was present could explain why the karyotypes from the two species did not initially cluster as expected. For 2La, we are able to accurately resolve karyotypes for the *An. gambiae* samples, but we are not able to analyze the *An. coluzzii* samples as only one sample had a different karyotype.

Our framework described here enables karotyping of inversions that had not been experimentally assessed. For example, by analyzing the 150 Burkina Faso *Anopheles* samples separately by species, we found potential 2Rc inverted regions in *An. coluzzii* (but not *An. gambiae*). Although the 34 *Anopheles* samples were not karyotyped for inversions on the 2R chromosome arm, we identify the potential presence of the 2Rj, and 2Rcu or 2Rbk inversions systems and their association with samples from Mali. Likewise, we confirm the presence of the 2Rb inversion in the 34 original *Anopheles* samples, which is expected given its presence in the Burkina Faso *Anopheles* samples.

In summary, not all PCs identify inversions when confounding factors are present. This will affect methods based purely on cluster structure in PCA projection (e.g., Ma et al. (2014); Cáceres and González (2015))); by using association tests and Manhattan plots, our proposed framework is able to distinguish between PCs capturing inversions versus others. This is expected given the role of PCA in population inference and other tasks (Lee et al. (2009); Patterson et al. (2006); Price et al. (2006); Neafsey et al. (2010)). It also is somewhat expected given prior modifications to augment PCA-based inversion detection. For example, Cáceres, et al. also analyzed linkage disequilibrium (Cáceres and González (2015); Sindi and Raphael (2010); Cáceres et al. (2012)) to better localize the inversions predicted by their likelihood-ratio framework, which assumes there will be three PCA clusters. Real-world data, however, violate typical assumptions due to confounding factors (species differences, more than three or muddled clusters) or unobserved karyotypes (two clusters instead of three), and we provide concrete examples for future evaluation of SNP-based inversion detection. In cases where the choice of PCs and number of clusters is ambiguous, the visualization of the associations provided by our cluster-SNP association approach can guide the required choices, which we show using inversions on the 2R arm in the *Anopheles* samples. Further, if karyotyping is not needed, our approach based on PC-SNP association tests eliminates the requirement of clustering completely.

## CONCLUSIONS

PCA-based frameworks can be used to detect, localize and karyotype inversions using only SNPs. We assess these approaches using data that varied in complexity from a single inversion in simulated samples to real sequencing data with multiple overlapping inversions, generated from multiple species and multiple geographic locations. Although we detect inversions on 2R in *Anopheles* data that had not been previously annotated, our analysis also confirms that PCA-based clustering can be affected by confounding factors, of which we present two actual manifestations for future SNP-based inversion detection assessment.

## ACKNOWLEDGMENTS

We would like to thank Michelle Riehle, Katrina Schlum, Jenica L. Abrudan, Christopher Beal, Derek Riley, and Josiah Yoder for insightful discussions and feedback that have improved our study and manuscript. RJN and KRM gratefully acknowledge MSOE for funding.

